## Supplementary Figure 1 for "Recognition of BRAF by CDC37 and Re-Evaluation of the Activation Mechanism for the Class 2 BRAF-L597R Mutant" (from another version of this preprint: version 1, 2022-05-20)

| Cdc37 construct | Cdc37 Structure | Binding partner | Binding parameters | Refer to Fig |
| --- | --- | --- | --- | --- |
| Full-length | 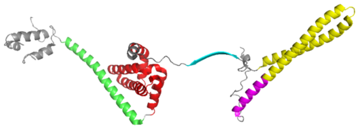 | sBraf | *Kd* = 1.0 μM | 1A |
| Full-length | 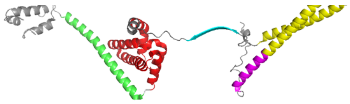 | sBrafV600E | *Kd* = 0.41 μM. | 1B |
| Full-length | 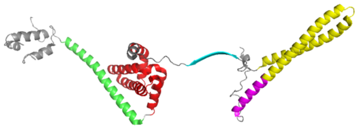 | sBrafV600E+ AMPPNP | NB | 1C |
| Full-length | 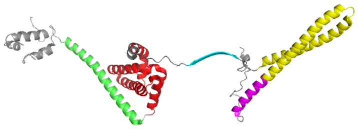 | sBrafV600E+ ADP | NB | 1D |
| N 1-120 | 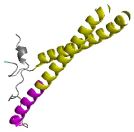 | sBrafV600E | *Kd* = 278 μM | 2B |
| C-terminal Domain 273-348 | 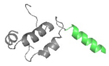 | sBrafV600E | *Kd* = 104 μM. | 2C |
| Middle domain 148-269 | 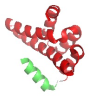 | sBrafV600E | NB | 2D |
| Cdc37 TrpZip | 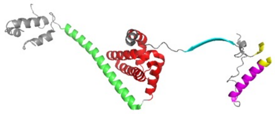 | sBrafV600E | *Kd* = 0.5 μM. | 3A |
| Cdc37 Δ(CC-βS) | 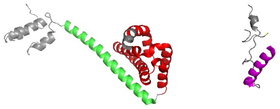 | sBrafV600E | *Kd* = 0.61 μM | 3B |

**Supplementary Figure 1 continued**

| Cdc37 construct | Cdc37 Structure | Binding partner | Binding parameters | Refer to Fig |
|---|---|---|---|---|
| Full-length | 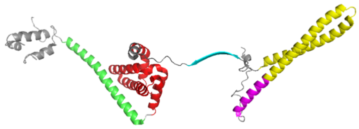 | sBrafV600E | *K*d = 0.41 μM. | 1B |
| Cdc37 Δ(7-CC-βS-272-285) | 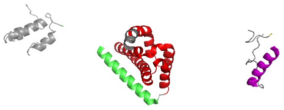 | sBrafV600E | *K*d = 1.03 μM. | 3C |
| Cdc37 $N^m$C | 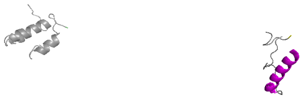 | sBrafV600E | *K*d = 145 μM. | 3D |
| Cdc37 $N^m$(+14)-C | 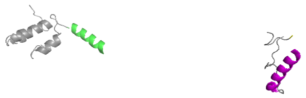 | sBrafV600E | *K*d = 8.7 μM | 4A |
| Cdc37 $N^m$(+6)-C | 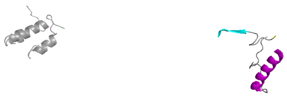 | sBrafV600E | *K*d = 18 μM | 4B |
| Cdc37 $N^m$(+41)-C | 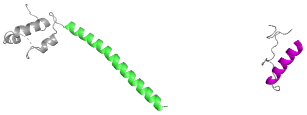 | sBrafV600E | *K*d = 1.99 μM | 4C |
| Cdc37 1-343 | 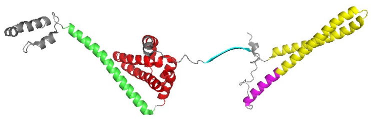 | sBrafV600E | *K*d = 0.76 μM | 5A |
| Cdc37 V343A | 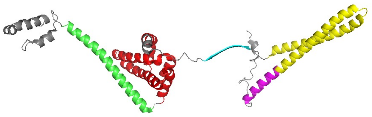 | sBrafV600E | *K*d = 0.9 μM. | 5B |
| Cdc37 V343R | 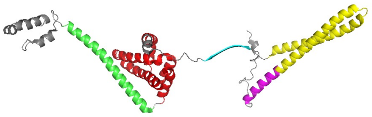 | sBrafV600E | *K*d = 1.3 μM. | 5C |

### Supplementary Figure 1 continued

| Cdc37 construct | Cdc37 Structure | Binding partner | Binding parameters | Refer to Fig |
|---|---|---|---|---|
| Full-length | 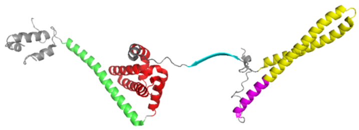 | sBrafV600E | *K*d = 0.41 μM | 1B |
| Cdc37 1-342 | 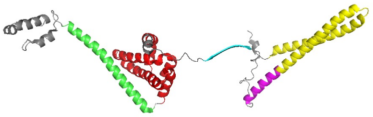 | sBrafV600E | *K*d = 45.7 μM | 5D |
| Cdc37 W342A | 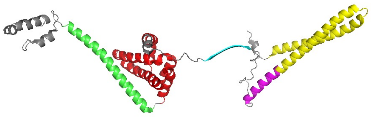 | sBrafV600E | *K*d = 94.3 μM | 5E |
| Cdc37 W342R | 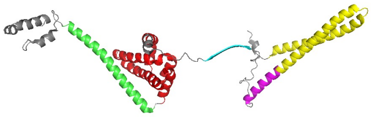 | sBrafV600E | NB | 5F |
| Full-length S13E | 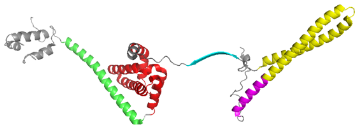 | sBrafV600E | *K*d = 0.30 μM | 6A |
| Cdc37 N22A | 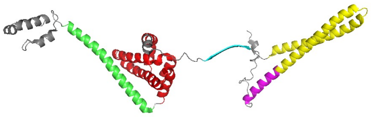 | sBrafV600E | *K*d = 27.3 μM | 7A |
| Cdc37 N22R | 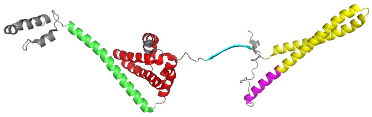 | sBrafV600E | *K*d = 14.2 μM | 7B |

**Supplementary Figure 1 continued**

| Cdc37 construct | Cdc37 Structure | Binding partner | Binding parameters | Refer to Fig |
|---|---|---|---|---|
| Full-length | 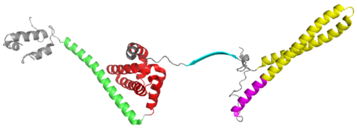 | sBrafV600E | *K*d = 0.41 μM | 1B |
| Full-length L28R | 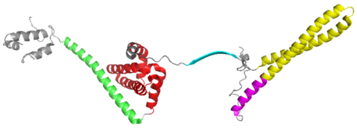 | sBrafV600E | NB | 8B |
| Full-length L28A | 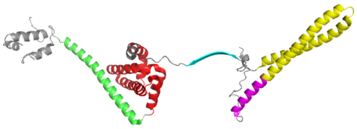 | sBrafV600E | *K*d = 16.9 μM. | 8C |
| Full-length W31K | 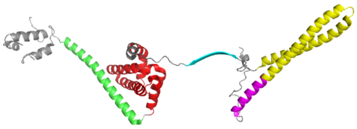 | sBrafV600E | NB | 8D |
| Full-length W31A | 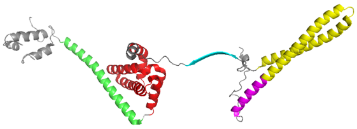 | sBrafV600E | *K*d = 2.1 μM | 8E |
| Full-length S27K | 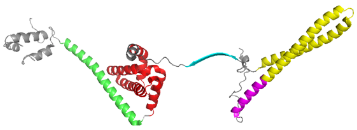 | sBrafV600E | NB | 8F |
| Full-length S27A | 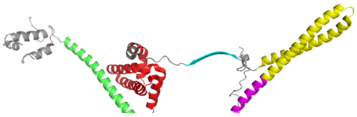 | sBrafV600E | *K*d = 1.6 μM. | 8G |
| Full-length Q47R | 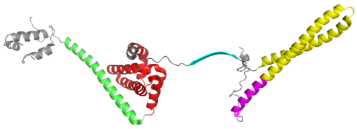 | sBrafV600E | *K*d = 0.3 μM | 8H |
| Full-length Q347A | 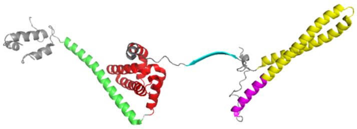 | sBrafV600E | *K*d = 0.35 μM. | 8I |

Supplementary Figure 1 continued

| Cdc37 construct | Cdc37 Structure | Binding partner | Binding parameters | Refer to Fig |
|---|---|---|---|---|
| Full-length | 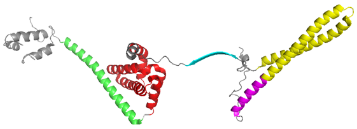 | sBrafV600E | *Kd* = 0.41 μM | 1B |
| Full-length | 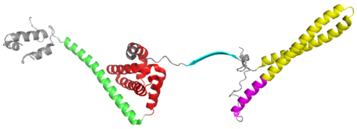 | sBrafV600E T599E-V600E-S602D | *Kd* = 0.47 μM | 9A |
| Full-length | 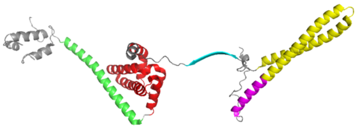 | sBrafV600E T599W-V600E | *Kd* = 0.23 μM | 9B |
| Full-length | 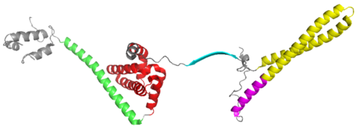 | sBraf F595A | *Kd* = 0.5 μM | 10A |
| Full-length | 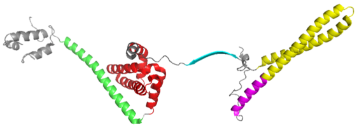 | sBraf F595R | *Kd* = 1.4 μM | 10B |
| Full-length | 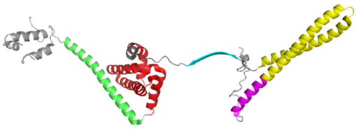 | sBraf D594V | *Kd* = 1.2 μM | 10C |
| Full-length | 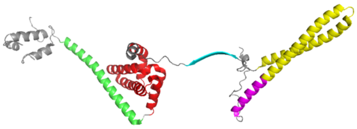 | sBraf L597A | *Kd* = 0.23 μM | 10D |
| Full-length | 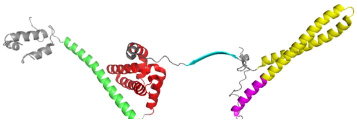 | sBraf L597R | NB | 10E |
| Full-length | 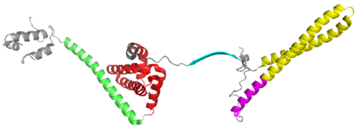 | sBraf L597R-V600E | *Kd* = 0.34 μM | 11D |

**Supplementary Figure 1 Constructs used in this study.**

**Refer to Fig 2A for amino acid positions of the various structural elements of Cdc37. *K*d values and the figure that each experiment represents is indicated.**
